# Regional Variant Analysis of Spike Glycoprotein Mutations of SARS-CoV-2 and Its Implications in COVID-19 Pandemic Control

**DOI:** 10.1101/2021.08.02.454771

**Authors:** Punnoth Poonkuzhi Naseef, Mohamed Saheer Kuruniyan, Shyju Ollakkod, U.K. Ilyas, Muhammed Elayadeth-Meethal

**Affiliations:** Department of Pharmaceutics, Moulana College of Pharmacy, Perinthalmanna, Kerala, 679321, India.; Dept. of Dental Technology. COAMS. King Khalid University. Abha. Saudi Arabia.61421.; Department of Animal Breeding and Genetics, College of Veterinary and Animal Sciences, Kerala Veterinary and Animal Sciences University, Wayanad, Kerala, 673576, India.; Department of Pharmacognosy and Phytochemistry, SRF, Faculty of Pharmacy, Hamdard University, New Delhi, India, 110062.; Department of Animal Breeding and Genetics, College of Veterinary and Animal Sciences, Kerala Veterinary and Animal Sciences University, Pookode, Wayanad, Kerala, India.

**Keywords:** SARS-CoV-2, COVID-19, Next generation sequence analysis, Virus-Host interaction, Spike glycoprotein, Mutation, Network analysis

## Abstract

Mutations in the spike glycoprotein have various impacts on the receptor binding, antibody interaction, and host range of SARS-CoV-2. As the interaction of spike glycoprotein with the human ACE2 receptor is the entry point of SARS-CoV-2 in human cells, mutations in the spike protein itself contain numerous impacts on the pandemic. Here, we analysed all the mutations in the spike glycoprotein from123 strains isolated from Kerala, India. We also predicted the possible structural relevance of the unique mutations based on topological analysis of the residue interaction network of the spike glycoprotein structure.

## 1. Introduction

The COVID-19 or coronavirus disease-19 caused by the severe acute respiratory syndrome coronavirus 2 (SARS-CoV-2) is an ongoing worldwide pandemic [1]. Although the original source of this virus to humans is still unknown, it first appeared in Wuhan, Hubei, China, in late 2019 [2]. In India, the first case of COVID-19 was reported in Thrissur, Kasargode, and Alappuzha, in Kerala, from where three medical students returned from Wuhan, China [3].

On 23 March 2020, the 1st lockdown was announced in Kerala. On 12 May 2021, Kerala was announced as the largest single-day state with 43529 new cases. On 24 June 2021, 2854325 are confirmed cases, a positive test case rate of 10.37%, and 2741436 recoveries and 12581 deaths in Kerala [4]. As of May 2021, Kerala has the 2nd highest number of confirmed cases of covid 19 after Maharashtra in India. In May 2021, more than 90% of known cases were reported due to community spread. The most affected districts are Ernakulam (12.2%), Malappuram (11.5%), and Kozhikode (10.6%) [5,6].

The RNA genome of coronavirus is covered by a helical capsid made up of N (nucleocapsid) proteins [7]. Due to its enveloped nature, the nucleocapsid is surrounded by M (membrane), E (envelope), and S (spike) protein [8]. In this viral structure, spike protein is especially important and draws research attention as the viral entry inside the host cell is mediated by this protein. Spike protein, which is a class 1 membrane fusion protein, has a transmembrane domain which helps to anchor the envelope [9]. The ectodomain radiates out from the viral structure which is solely responsible for receptor binding (via the S1 subunit) and membrane fusion (via the S2 subunit). S protein helps to fuse the viral and host membrane by changing its conformation. SARS-CoV interacts with the host through a hinge-like movement of the S1 subunit in the prefusion state which helps the S2 subunit into a dumbbell-shaped stable post-fusion conformation. This S2 subunit with its 6 helices brings the host and viral membrane together. S1 subunit has the receptor-binding domain for human ACE2 receptors as reported by various studies. Various mutations along the spike protein affect not only the receptor affinity but also the host range [10].

In the present study, we identified a total of 298 mutations in the spike protein from 123 SARS-CoV-2 isolates of Kerala, India. For the known mutations, we have reviewed their consequences in the SARS-CoV-2 structure, host range, and antibody binding capacity. But, for unknown mutations, which are unique to Kerala, we have tried to predict the mutational consequences using the residue interaction network. Here, the residue interaction network represents a complex network representation of protein structure where amino acids are nodes. We calculated network matrices and based on topological significance we predicted the possible effect of the mutations on the protein structure.

## 2. Results

We identified a total number of 298 spike mutations in various SARS-CoV-2 variants isolated from Kerala, India. In figure 1 we have included a pie chart of the various spike mutations with their abundance frequency.

**Figure 1.**
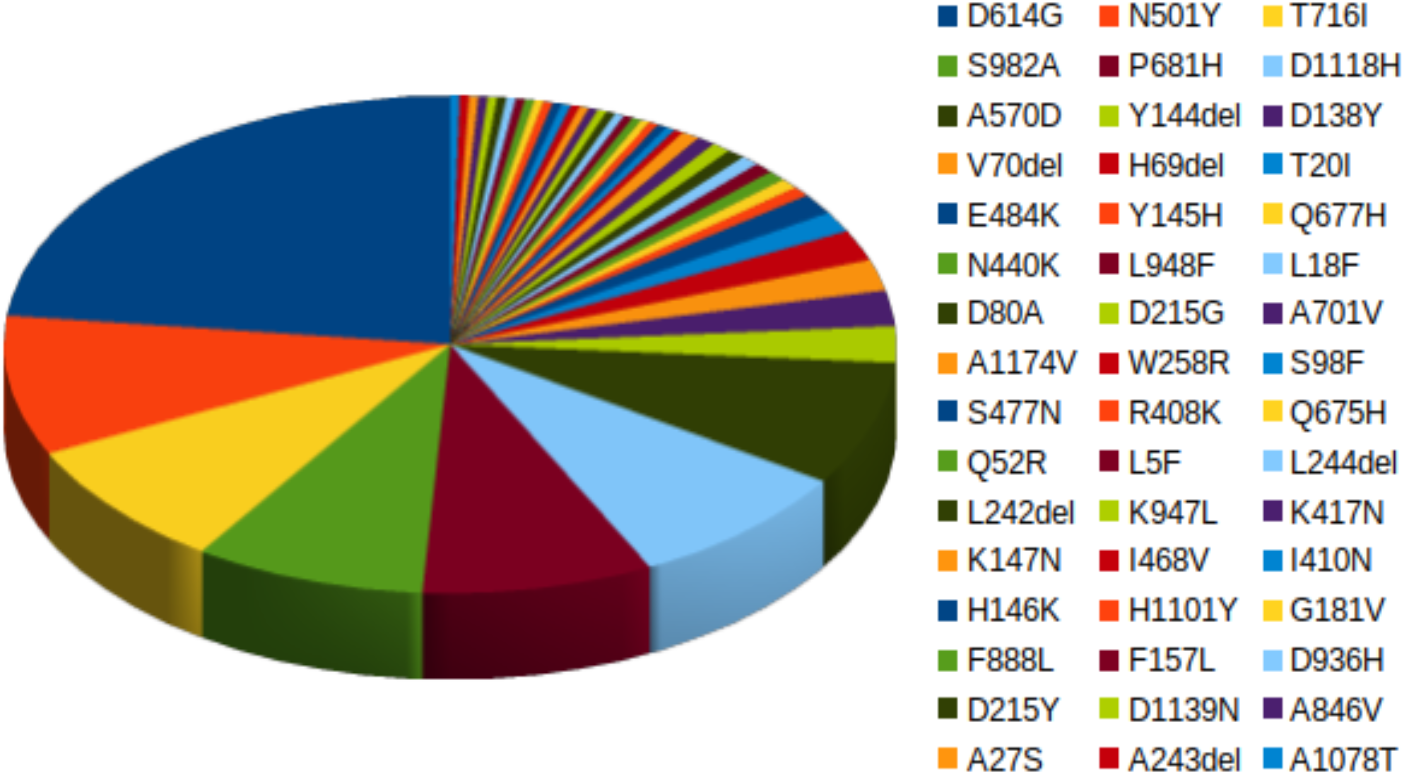
Frequency of 298 spike mutations in various SARS-CoV-2 variants isolated from Kerala, India

### RIN network analysis

To predict the effect of unique mutations on the protein structure, we have used topological analysis of the residue interaction network analysis based on node metrics. We have used spike glycoprotein of SARS-CoV (PDB id: 6ACC) for our protein residue interaction network construction [15]. We measured the change in node betweenness centrality metrics of every amino acid as the effect of the deletion of the mutated amino acids from the RIN. In figure 2, we represent the network view (a) of the spike glycoprotein with comparison to its three-dimensional helical structure (b).

**Fig 2.**
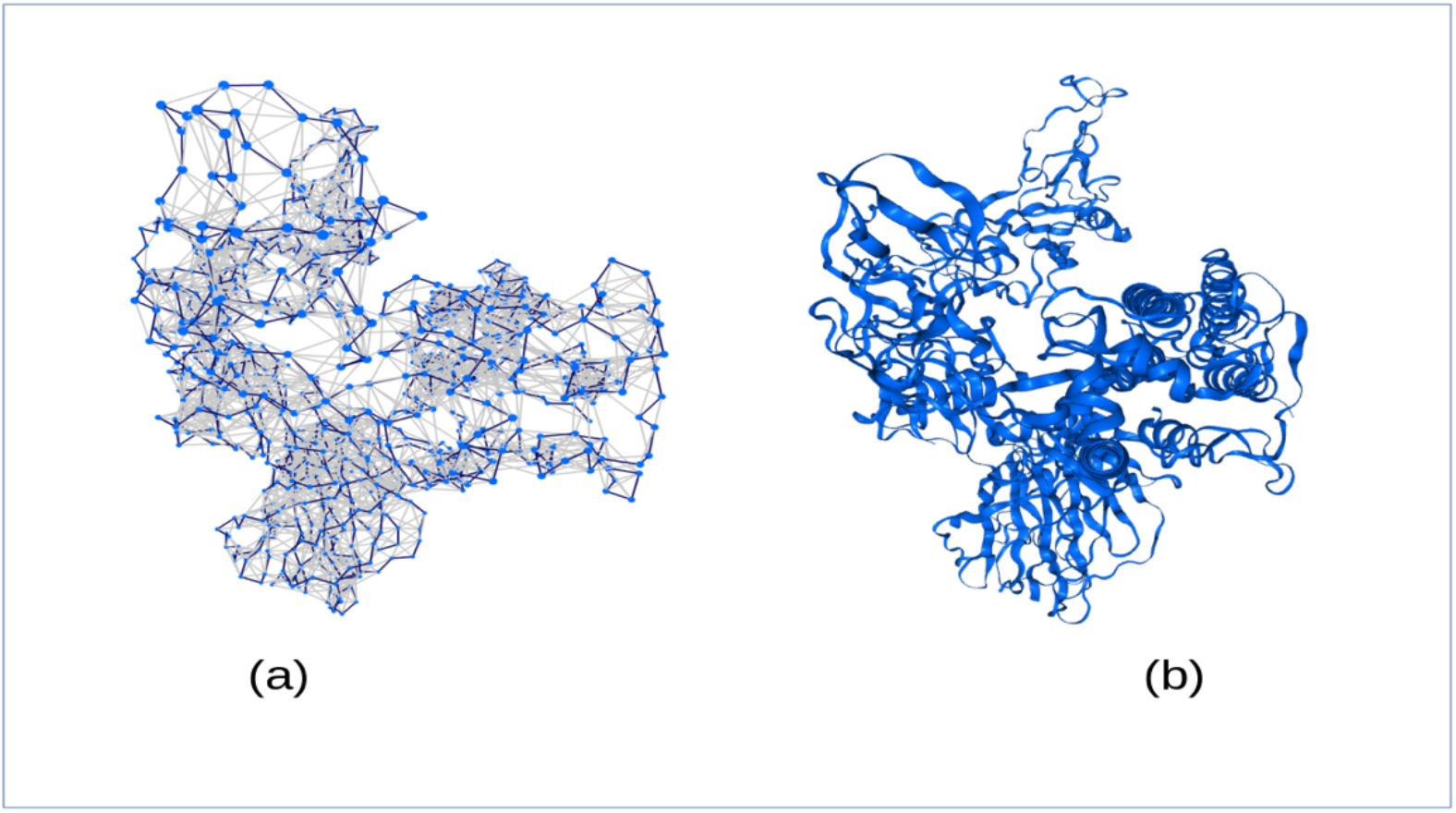
Network representation of spike glycoprotein in comparison to its three-dimensional helical structure

We measured the node centrality values for all the mutated amino acids which are unique to Kerala as represented in table 1.

| Node | Abundance | Degree | Z_D | Closeness | Z_C | Betweenness | Z_B |
| --- | --- | --- | --- | --- | --- | --- | --- |
| D614G | 69 | 8 | 0.25 | 0.062434 | 1.11 | 0.058986 | 0.21 |
| N501Y | 27 | 9 | 0.7 | 0.050503 | -0.24 | 0.001544 | -0.33 |
| T716I | 25 | 6 | -0.65 | 0.050681 | -0.22 | 0.003744 | -0.31 |
| S982A | 25 | 11 | 1.59 | 0.047371 | -0.59 | 0.008894 | -0.26 |
| P681H | 25 | 6 | -0.65 | 0.045263 | -0.83 | 0.001228 | -0.33 |
| D1118H | 25 | 7 | -0.2 | 0.045317 | -0.82 | 0.000634 | -0.33 |
| A570D | 24 | 6 | -0.65 | 0.055296 | 0.3 | 0.017623 | -0.18 |
| Y144del | 7 | 4 | -1.54 | 0.064724 | 1.36 | 0.005417 | -0.29 |
| D138Y | 7 | 4 | -1.54 | 0.066475 | 1.56 | 0.99097 | 8.89 |
| V70del | 6 | 6 | -0.65 | 0.049148 | -0.39 | 0.001561 | -0.33 |
| H69del | 6 | 10 | 1.15 | 0.054494 | 0.21 | 0.063394 | 0.25 |
| T20I | 4 | 9 | 0.7 | 0.055047 | 0.28 | 0.001147 | -0.33 |
| E484K | 4 | 9 | 0.7 | 0.058918 | 0.71 | 0.019155 | -0.16 |
| Y145H |  | 7 | -0.2 | 0.054106 | 0.17 | 0.002706 | -0.32 |

It was observed that the mutation D138Y could be very crucial as the position has a very high betweenness value. It indicates that the amino acid of position 138 is very important as many shortest paths pass through this point. Mutation at this point changes the interaction which alters the overall structure of the protein. Residues with high betweenness indicate their importance in ligand binding [16]. This position also has a very high closeness value which indicates that mutation in this position also could affect various ligand or receptor binding as indicated in the previous literature [17]. Other than that mutation Y144del and D614G are also important based on their closeness values. Mutation in S982A and H69del has a very high degree which indicates that the amino acids in these two positions interact with many other amino acids in the protein structure. So, the mutation in these two positions also should have biological significance.

## 3. Discussion

In this paper, we have summarised all the 298-point mutations in the spike glycoprotein which are isolated from Kerala, India [7,8]. Among them, we identified the top 14 mutations based on their abundance in the SARS-CoV-2 variants of Kerala. Further, we summarised the abundance frequency of these mutations in Kerala. To explore their biological significance, we used residue interaction network analysis. Based on some well-known topological measures we predicted the significant mutations among these 14 unique mutations.

Our network analysis result is based on node degree, closeness, and betweenness centrality. In a previous study, a residue interaction network study found that more than 1/3 of biologically essential residues have been identified to possess high closeness values [15]. Not only that, residues in the active sites and that are associated with ligand binding activity, also possess high closeness values. Betweenness centrality also indicates the ligand-binding sites as reported previously [5]. So, our prediction of the potential biological significance of these unique spike mutations needs to be studied further with experimental verifications. We believe this research will help to understand the SARS-CoV-2 spike mutations and would help researchers in the development of alternative vaccines.

## 4. Materials and Methods

### Identification of spike mutations

The study was based on a total number of 123 SARS-CoV-2 full genomes of Kerala origin fetched from the GISAID database [11]. Multiple sequence alignment was performed with the reference sequence of hCoV-19/Wuhan/WIV04/2019, which has been isolated from Wuhan of China and has been considered the reference strain worldwide [1]. With the help of the CoVsurver mutation analysis tool of SARS-CoV-2, we have identified all the mutations on the spike protein of SARS-CoV-2 isolates from Kerala, India [12].

### Residue interaction network (RIN) construction

For the construction of a residue interaction network (Brinda et al. 2005) the respective PDB file of the protein is required [13]. Residue interaction network depends on the three-dimensional coordinates of each atom of amino acid. Here, we have mainly focused on the C-alpha network which is based on the C-alpha atom of amino acid. In this network each of the C-alpha atoms of each amino acid is considered as the node representing the amino acid in the network. An edge will be considered between two amino acids if the cutoff distance between the two respective C-alpha atoms is >= 7Å. For the construction of the network, we have used NAPS web-server [14].

### Network metric analysis

We calculated the basic network metric for each mutated node of the network. Each of the network metrics has its biological significance as reported in various literature [18]. In a graph, a node’s degree represents the number of connections the node has with other nodes in the network. Closeness centrality is defined by the reciprocal of the shortest path distance of that node to every other node in the graph. So, the closeness centrality of a node i in a network is represented by the following formula,

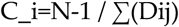

Where N is the total number of nodes in the network. Dij represents the shortest path distance between i and j in the network. Now the closeness centrality of a node in a biological network could reveal various importance.

Similarly, the betweenness centrality of a node represents the number of shortest paths passing through that node. Betweenness centrality of node v is represented as,

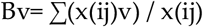

Where, x(ij) is the number of the shortest paths from node i to j in the network and x(ij)v is the number of the shortest paths from node i to j passing through v.

Also, to compare a node’s topological metric in respect to the same of other nodes, we have calculated the Z score values. The Z score of a measure of a node is defined as,

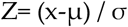

where x is the value of the measure of the node. μ and σ represent the mean and standard deviation of the same measure of the whole population. We have considered only Z value >=1.

## Author Contributions

Conceptualization, NPP, MSK, MEM.; methodology, NPP, MEM.; software, NPP, MSK, MEM.; data curation, NPP, MEM, UKI.; writing–original draft preparation, NPP, MEM; writing–review and editing, NPP, MEM, MSK, UKI, SO.; supervision, MEM.; project administration, NPP, MEM, MSK.

## Acknowledgments

The authors extend their appreciation to the Deanship of Scientific Research at King Khalid University, Saudi Arabia for funding this work through the Research Group Program under Grant No: RGP 2/ 191/42.

